# Two-zone tumor tolerance can arise from a simple immunological feedforward motif that estimates tumor growth rates

**DOI:** 10.1101/095455

**Authors:** Eduardo D. Sontag

## Abstract

Since the early 1990s, many authors have independently suggested that self/nonself recognition by the immune system might be modulated by the rates of change of antigen challenges. This paper introduces an extremely simple and purely conceptual mathematical model that allows dynamic discrimination of immune challenges. The main component of the model is a motif which is ubiquitous in systems biology, the incoherent feedforward loop, which endows the system with the capability to estimate exponential growth exponents, a prediction which is consistent with experimental work showing that exponentially increasing antigen stimulation is a determinant of immune reactivity. Combined with a bistable system and a simple feedback repression mechanism, an interesting phenomenon emerges as a tumor growth rate increases: elimination, tolerance (tumor growth), again elimination, and finally a second zone of tolerance (tumor escape). This prediction from our model is analogous to the “two-zone tumor tolerance” phenomenon experimentally validated since the mid 1970s. Moreover, we provide a plausible biological instantiation of our circuit using combinations of regulatory and effector T cells.

## 1 Introduction

In vertebrates, the innate and adaptive immune systems combine to provide a finely orchestrated multicomponent host defense mechanism, which protects against infective pathogens (viruses, harmful bacteria, and other microorganisms) and also helps to identify and eliminate malignantly transformed cells (immune surveillance of tumors) [33, 1]. A prerequisite for successful immune action is the ability to distinguish foreign agents from an organism’s own healthy tissue (self/nonself dichotomy). A central role in this discriminatory ability is played by antigens, which are molecules capable of inducing an immune response. Nonself antigens appear not only in pathogens, but also in malignantly transformed cells due to their overexpression of normal proteins, mutated proteins, or oncogenic viruses. In its normal functioning, the immune system is trained, through a variety of means such as the negative selection of T cells in the thymus, to not react to self-antigens. This paradigm of self/nonself pattern recognition traces back to Burnet [8], and related ideas in [62].

The static view, however, is not entirely consistent with a number of phenomena which hint to a role for discrimination based on dynamic features, as suggested by the following examples: (1) The presence of commensal bacteria (microbiome) is stably tolerated by the immune system, despite the presence of bacterially derived, non-self molecules that exist in intimate proximity to the host [49]. (2) Slow-growing tumors are known to evade the immune system [23] despite their expression of non-self antigens. (3) Decreased activation of natural killer cells is observed under chronic receptor activation [48] despite the continued presence of antigens. (4) Endotoxin tolerance in macrophages, also known as deactivation, desensitization, adaptation, or reprogramming, refers to the reduced capacity of a host to respond to the pro-inflammatory stimuli of bacterial signatures such as lipopolysaccharides after a first exposure to the same type of stimulus [68], thus helping to avoid harm caused by their continual presence. (5) The phenomenon of anergy, in which lymphocytes such as B and T cells fail to respond to their specific antigen, might suggest additional regulatory mechanisms [24]. On the other hand, it could be argued that autoimmune diseases contradict this dynamic view, since in that case the immune system attacks basically constant antigen loads. However, as Pradeu and collaborators argue [49], many autoimmune diseases appear after relatively fast changes occurring during puberty or due to a sudden exposure to chemical or biological agents and, conversely, allergy treatment by slow desensitization through antigen exposure leads to tolerance [7]. These shortcomings suggest that one might want to also consider *dynamic* features of antigen presentation as a complement to discrimination mechanisms which are only based on a static self/nonself dichotomy.

During the past 30 or so years, a number of authors, most notably Zvi Grossman and collaborators [23] and Pradeu and collaborators [49], have proposed the necessity of incorporating dynamics in self/nonself recognition. They support this line of reasoning by experimental observations that (a) lymphocytes mount a sustained response only when faced with a sufficiently steep increase in their level of stimulation (antigen presentation, proliferation rates of infected cells or tumors, stress signals) and, dually, (b) even when a new motif triggers an immune response, its chronic presence may result in adaptation: downregulation or even complete termination of the inflammatory response. One mathematical formulation was introduced by Grossman and Paul [24], who in 1992 postulated the “tunable activation threshold” model for immune responses: effector cells in the innate or adaptive systems should become tolerant to continuously expressed motifs, or even gradually increasing ones, but should induce an effector response when a steep change is detected. Among recent variations upon this theme are the “discontinuity theory” postulated by Pradeu, Jaeger, and Vivier [48], and the “growth threshold conjecture” due to Arias, Herrero, Cuesta, Acosta, and Fernández-Arias [4]. In Section 5 we briefly review these models.

### 1.1 A simple model

In that context, we propose and analyze here a novel and simple mathematical model, consisting of three ordinary differential equations as follows:

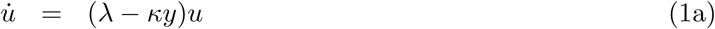

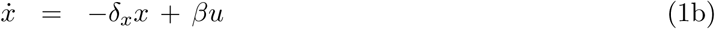

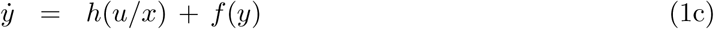

(dots indicate time derivatives), see Figure 1. The constants λ, κ, δ_x_, β are positive. The function *h* is continuous, strictly increasing, and satisfies *h*(0) = 0; for our purposes we could simply take a linear function, *h*(*p*) = *μp*. Qualitatively, the plot of the function *f* is as in the left panel of Figure 1(B).

**Figure 1:**
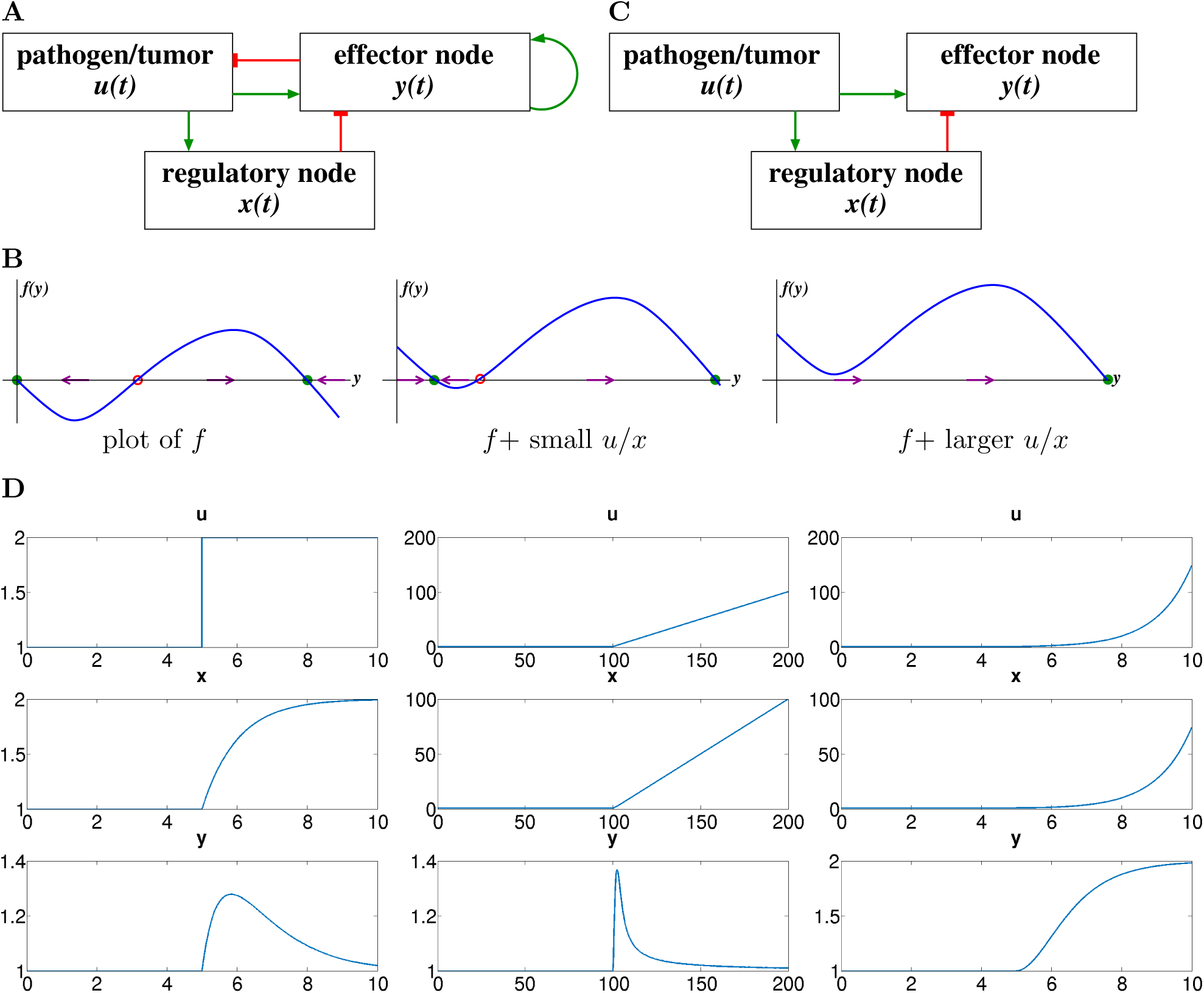
A. The model being considered in this paper. Blunt-end red arrows denote repression, green arrows denote activation or autocatalytic feedback. B. Plot of the nonlinear function *f* that endows the system with bistability (left), and two translates of this plot; open red and filled green circles denote unstable and stable states respectively. C. Simplified model when feedbacks are ignored. D. Simulations of system (2), with input *u*(*t*) switching in the middle of the interval from *u*(*t*) = 1 to a new constant value *u*(*t*) = 2, a shifted ramp *u*(*t*) = *t*, and a shifted exponential *u*(*t*) = *e^t^* respectively; also shown are *x*(*t*) and *y*(*t*). In every case, the horizontal axis is time, *t*. Parameters used: *δ_x_ = β = μ = δ_y_* = 1; initial conditions *x*(0) = *y*(0) = 1.

We view the system described by equations (1) as a “toy model” that encompasses immune suppression as well as pattern discrimination based on antigen dynamics. The variable *u* represents an immune challenge, such as the volume or number of cells in a tumor, an infection, or the amount of antigen-presenting cells (APC) of a certain type, and the term λ*u* in the equation for *u* represents its exponential rate of growth. The variable *x* represents an intermediate regulatory node, such as the amount of T regulatory cells in a tumor microenvironment, and this variable evolves according to a linear activation, proportional to the immune challenge, and decays linearly (inactivation or degradation). The variable *y* represents an agent that can eliminate the challenge *u*, such as the number of tumor-specific cytotoxic T cells (CTL) in the same environment. This elimination is represented by the mass-action term *—κyu* in the equation for *u*. The variable *y* increases in proportion to the ratio of *u* to *x*, which implies that *y* is driven by the rate of growth of *u*, as will be explained later. The function *f* (*y*) combines degradation and a positive autocatalytic feedback such as induced by cytokines, and is chosen so as to endow the *y* component with bistable behavior, as explained next.

### 1.2 Intuitive discussion of model

Initial intuition regarding the system (1) can be obtained by thinking of *p* = *h*(*u/x*) as a parameter in the scalar differential equation (1c), writing *ẏ* = *p* + *f*(*y*) and temporarily ignoring that in the full model *p* is not constant but it depends on *y* through a feedback repression of *u*. Note that the plot of *p* + *f* (*y*) is a vertical translation by *p* of the plot of *f*. If *p* is small, then, starting from the initial condition *y*(0) = 0, the solution *y*(*t*) approaches asymptotically a low value of *y*, the left-most stable green-labeled point in the center plot in Figure 1(B). However, once that *p* has a value large enough that this first equilibrium disappears (in what is known as a “saddle-node bifurcation”) then *y*(*t*) will converge, instead, to a comparatively large value, the green stable point in Figure 1(B), right. This higher equilibrium represents the triggering of a substantially increased immune response.

Further intuition can be gleaned from another simplification. Let us think of *u* as an input to equations (1b,c), once again ignoring the repression of *u* by *y*, see Figure 1(C). In addition, let us take *h*(*u*/*x*) = *μ u/x* and let the function *f* include only degradation or deactivation but no autocatalytic feedback. Thus (1b,c) simplify to:

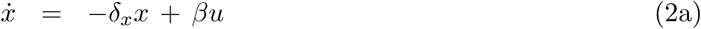

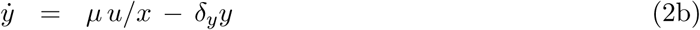

where *β, μ, δ_x_,δ_y_*, are some positive constants and *u* is viewed as an external stimulus. The resulting system is a type of incoherent feedforward loop (IFFL), a “motif” which is ubiquitous in biological networks [45, 2], and is statistically enriched in many intracellular networks (metabolic pathways, genetic circuits, kinase-mediated signaling) as well as the intercelullar level. IFFL’s are characterized by the existence of two antagonistic (“incoherent”) alternative pathways from the input to the output. In a direct path, the external cue or stimulus *u*, let us say a step function as shown in the left panel of Figure 1(D), activates the effector node *y* as well as a regulatory node *x* which in turn represses *y*. This structure endows IFFL’s with powerful signal processing capabilities, studied in detail in Alon’s textbook [2]. As the indirect effect requires an accumulation of *x* over time, there is typically a delay in the downregulation of *y*, leading to a response that consists of a short activity burst (fold detection) followed by a return to a basal value which is the same no matter what was the magnitude of the input, as shown in the left plots in Figure 1(D). The return to a basal value independent of the excitation magnitude is a phenomenon called “perfect adaptation” [2, 56], and holds also for ramps (linearly increasing) inputs *u*(*t*), see center plots in Figure 1(D). However, when instead *u*(*t*) = *u*(0)*e^vt^* grows exponentially, *u̇* = *vu*, the response *y*(*t*) approaches a constant multiple of the rate *v*, see right plots in Figure 1(D). Thus, the system can be seen as an estimator of the growth constant *v*, and does not respond persistently to sub-exponential inputs. See [59] for a formal proof of this property. If we now take again into account the autocatalytic feedback term in *f*, then a large value of *u/x*, which as we just discussed depends on the steepness of the input *u*, may trigger an irreversible transition to a higher state for *y*, which persists even after the excitation goes away.

The term “fold detection” for the initial activity burst is motivated by the response to an input that switches from *u*(*t*) = *u*_−_ for *t* < 0 to *u*(*t*) = *u*_+_ for *t* ≥ 0: assuming that *x*(0) is adapted to *u*_−_, the initial amplitude of *u/x*, which triggers the initial change in response in *y*, is (*δ_x_/β*)*u*_+_*u*_−_ and *u_+_/u_−_* is the fold change in the input. In the IFFL system (2), an input that is scaled by a positive constant *p* leads to the same response *y*, provided that the regulatory variable *x* is also scaled by *p*, since one has 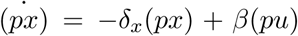 and *ẏ =μu/x − δ_y_y* = *μ*(*pu*)/(*px*) − *δ_y_*.

In mathematical terms, this says that the equations do not change under the one-parameter Lie group of transformations 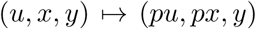. In other words, this system is “scale invariant” (or, in different terminology, it is a “fold change detector”) [52]. Section 7 includes a more detailed discussion of scale invariance for this system.

In summary, our model combines three central motifs in modern systems biology: (1) an IFFL for estimating the rate of growth of *u*(*t*), which is scale-invariant, (2) a multi-stable dynamics for *y*(*t*), and (3) feedback control of *y*(*t*) that represses the input *u*(*t*).

In spite of its extreme simplicity, analysis of the model (1) reveals a somewhat surprising phenomenon: there may exist more than one growth range for which an immune challenge, for example a tumor, can be eliminated. Specifically, we find three thresholds λ_1_, λ_2_, λ_3_, so that:

- if the per-capita rate of growth λ of the tumor is less than λ_1_, then it is eventually eliminated by the immune system;
- when the tumor is more aggressive, λ > λ_1_ but λ < λ_2_, it cannot be eliminated (it is “tolerated” by the immune system);
- for an even more aggressive challenge, with λ > λ_2_ but λ < λ_3_, again the tumor is eliminated; and
- if λ > λ_3_, again there is no elimination (the tumor “escapes”).

Intuitively, in the intermediate range λ_2_ < λ < λ_3_ the immune system goes into “overdrive,” engaging additional resources through activation of a positive feedback mechanism, and this level is powerful enough to effectively repress the challenge.

As we mentioned earlier, the impact of growth rates on immune responses has been a hot topic of discussion in the immunology literature, and it has been proposed that different rates provide a way to differentiate among threats based on their aggressivity. “Non-self” and potentially dangerous cells presumably reproduce faster, compared to “self” and also beneficial microorganisms. As we will discuss, the role of *exponential* rates in determining immune response has also been the subject of experimental research, including a recent immunotherapy patent, and the role of T suppressor cells in providing what we may now view as the regulatory node in an incoherent feedforward loop has been well-established experimentally as well.

The prediction of the existence of disjoint regions of tumor elimination, depending on rate of growth, remains to be tested. However, this behavior is strongly reminiscent of the well-known phenomenon of “two-zone” tumor tolerance, which has been observed in the experimental literature since the mid-1960s. This phenomenon is entirely analogous to our predictions, the only difference being the regions are now determined by the magnitude of an initial tumor inocula in animal subjects instead of growth rates. We discuss this and, more generally, the phenomenon of “sneaking-through” of tumors. Several other predictions also follow from the model, such as its capability for logarithmic sensing and scale invariance, each of which could be in principle experimentally tested. As an example of plausibility, we discuss how our model might represent the combined actions of T regulatory cells and cytokine feedback.

This preprint considerably extends the material in [58], but some material from that previous report is repeated here so as to make it self-contained. A version of this preprint has been submitted [60]. *However, the journal version does not include Section 9 on degradation-based IFFL’s, which has been now added.*

## 2 Results

We summarize now our main results

### A key reduction to a system of two equations

Scale invariance under the transformations 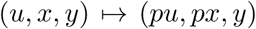 suggests performing a change of variables in which *x* is replaced by *p* = *u/x*. As *x* = *u/p*, the original variables (*u,x,y*) can be recovered from (*u, p, y*), so the transformation is invertible. Rearranging the order of equations, we will from now study the system (1) in these new coordinates:

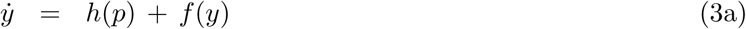

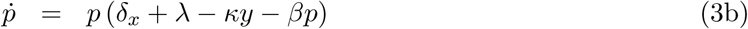

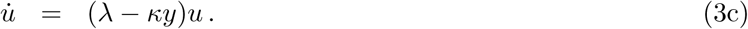

Writing the equations in these new coordinates has a major advantages for analysis, because the equations for *y* and *p* are now decoupled from *u*, and therefore we may study the two-dimensional (*y*,*p*) system using techniques suitable for planar systems, such as phase planes and nullclines. Information regarding the asymptotic behavior of *u* can be inferred from that of the (*y*, *p*) system. Suppose that we have determined that a solution (*y*(*t*),*p*(*t*)) tends to an equilibrium (*y̅*,*p̅*) with *p̅*≠0. This implies that λ−*κy*→λ−*κy*̅ as *t* → ∞, and, because *δ_x_+*λ−*κy*̅−*βp̅* = 0 at any equilibrium with *p̅*≠0, asymptotically *u*(*t*) ∝ *e^vt^* with *v=*λ*−κy̅−βp̅−δ_x_*

This allows us to decide if *u*(*t*) converges to zero or infinity (whether the immune challenge is eliminated or grows without limit):

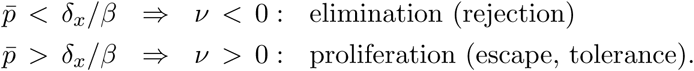

Thus, from now on we study (3a,b). Recall the plot of *f* is as in Figure 1(B), and that *h* is assumed to be strictly increasing with *h*(0) = 0, which we will later specialize to *h*(*p*) = *μp*. We are only interested in solutions with *y*(*t*) ≥ 0 and *p*(*t*) ≥ 0.

### Equilibria and nullclines of the reduced system

The equilibria of system (3a,b) are obtained by simultaneously solving

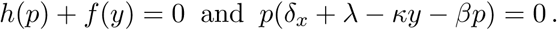

When *p* = 0 the first of these is simply *f* (*y*) = 0 (because *h*(0) = 0), and when *p* > 0, the second gives *δ_x_ + λ − κy − βp* = 0. The *y* and *p* nullclines of this system are the subsets of the first quadrant where *ẏ* = 0 and *ṗ* = 0, that is,

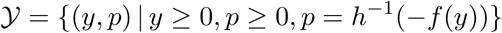

and

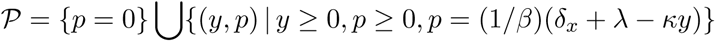

respectively. Note that (*y,p*) can only belong to *𝒴* if − *f*(*y*) is in the range of *h*, and in particular this means that *f*(*y*) ≤ 0, thus ruling out the values of *y* for which the plot of *f* is positive. Similarly, (*y,p*) can only belong to *𝒫* if *y* ≤ (*δ_x_ + λ*)/*κ*. The equilibria of the system are the points at the intersection of these two sets.

We are interested in analyzing the behavior of the system for different values of the parameter λ which quantifies the initial growth rate of the immune challenge. The only place where λ plays a role is the equation for the line *p* = (1/*β*)(*δ*_*x*_ + λ − *κy*), picking one of its parallel translates. See Figure 2.

**Figure 2:**
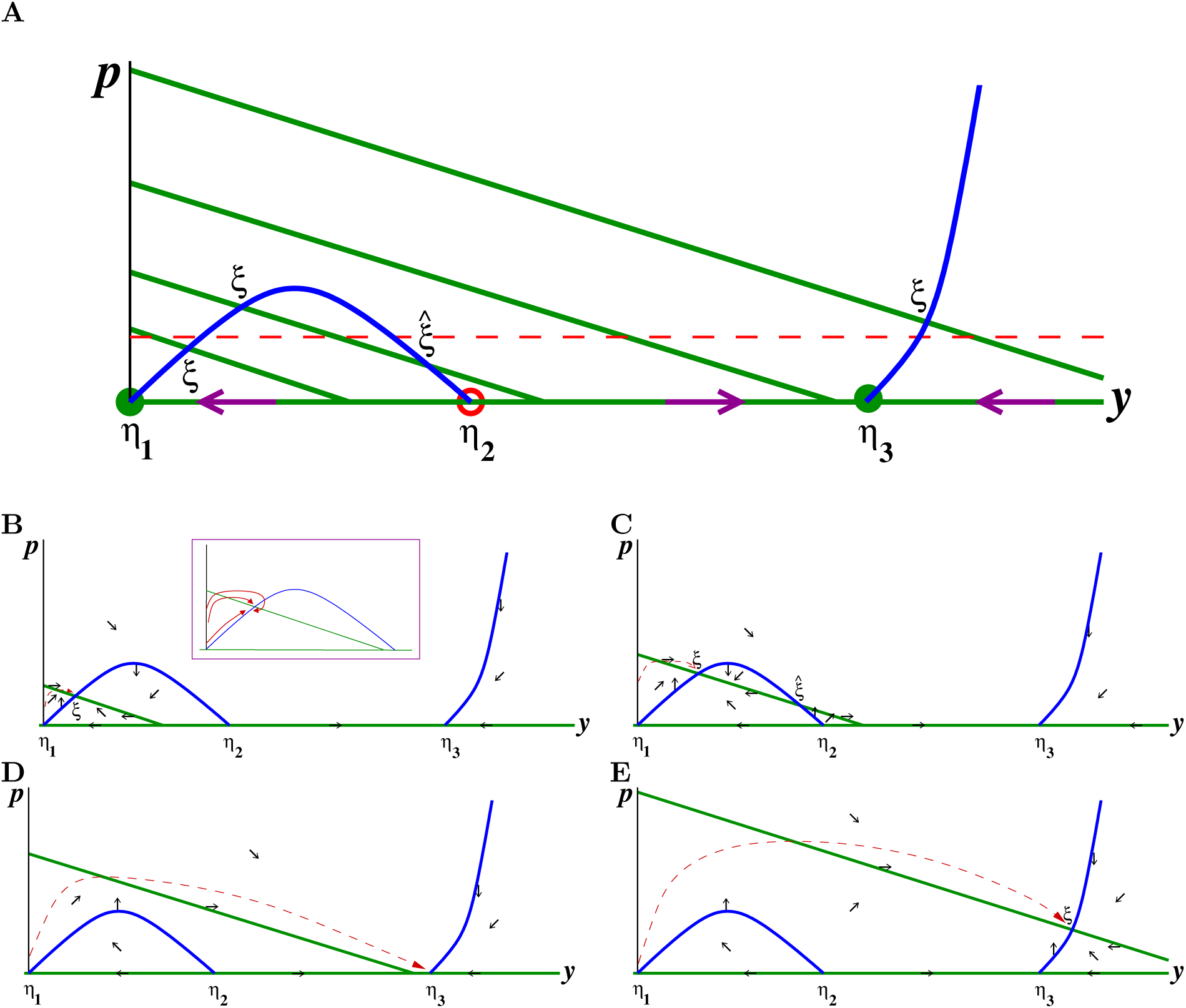
**A.** Nullclines for the system (3a,b) on the quadrant *y* ≥ 0, *p* ≥ 0. Blue curve is *y*-nullcline. Green lines are *p*-nullclines, shown for four typical values of λ. There are two different types of equilibria: the *η*’s are equilibria with *p* = 0 and the *ξ*’s with *p* > 0. The four points labeled are *ξ* stable for the reduced dynamics, for the respective λ’s, as discussed in the text. The intercepts of the green lines with the *p* axis (vertical) are at the locations *p* = (1/*β*)(*δ*_*x*_+ λ_*i*_), *i* = 1,2,3,4. Red dashed horizontal line is threshold *p = δ_x_/β* discussed in the text. **B-E.** Directions of movement in each region determined by the nullclines, along with typical trajectories, respectively, low to high for each of the values of λ represented in **A**. Inset in **B** shows various possibilities for approach of trajectory to point labeled *ξ*.

We assume, for simplicity, there are no more than two positive intersections between any (green) line *p* = (1/*β*)(*δ_x_ +* λ − *κy*) and the (blue) *y* nullcline; this means that the slope *− κ/β* is not ≈ 0. We also assume for the values of λ that we analyze that there is at most one intersection, like 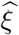, on the decreasing branch of the blue curve. (If there are such additional intersections, the theoretical analysis is somewhat more complicated, but is analogous.) The dashed line is the threshold that determines the behavior of *u*: if (*y*(*t*),*p*(*t*)) converges to an equilibrium that has *p̅ < δ_x_/β*, then *u*(*t*) → 0 as *t* → ∞, but if *p̅ > δ_x_/β*, then *u*(*t*) → ∞.

### Linearized stability at equilibria

The Jacobian matrix for the system (3a,b), evaluated at a generic equilibrium (*y*̅,*p*̅), is:

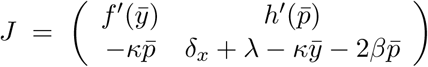

and therefore

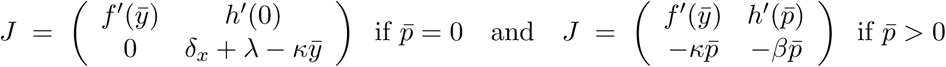

where we used that *δ*_x_ + λ − *κy* − 2*βp̅ = −βp̅* when *p*̅ > 0 (since the equilibrium condition is *δ_x_* + λ − *κy − βp̅* = 0 in that case). Note that these cases correspond, respectively, to points labeled *η* and *ξ* in Figure 2.

At points with *p*̅ = 0, the eigenvalues of the upper triangular matrix *J* are *f′*(*y*̅) and *δ_x_ +* λ − *κy̅*. Referring to the plot of *f* in Figure 1(B), *f′(y̅) >* 0 at the point *η*_2_ and *f′(y̅*) < 0 at the points *η*_1_ and *η*_3_. This is because the blue curve in Figure 2 is *h^−1^(−f*(*y*)) and therefore is qualitatively like an upside-down version of the plot of *f*, so that positive (respectively negative) *f′* corresponds to negative (respectively positive) slope at the equilibrium. Thus *η*_2_ is unstable. At the point *η*_1_, the second eigenvalue is *δ_x_ +* λ − *κy̅ = δ_x_* + λ, which is positive, so *η*_1_ is also unstable (a saddle point). The stability of *η*_3_ depends on the sign of *δ_x_ +* λ − *κy̅*: *η*_3_ is stable if this sign is negative, that is, if *δ_x_ +* λ − *κy̅* < 0, and unstable otherwise. Suppose that all parameters are fixed except for λ. We use Figure 2 to determine stability: if λ is such that the corresponding line *p = (1/β)(δ_x_ +* λ − *κy*) intersects the *y* axis at a point *y_0_* to the left of *η*_3_, that is, if *y*_0_ < *y̅*, then *δ_x_ +* λ − *κy_0_* = 0 implies *δ_x_ +* λ − *κy̅* < 0, and hence *η*_3_ is stable. If, on the other hand, λ is larger and *p = (1/β)(δ_χ_ +* λ − *κy*) intersects the *y* axis at a point *y*_0_ to the right of *η*_3_, then *δ_x_ +* λ − *κy_0_ =* 0 implies *δ_x_ +* λ − *κy̅* < 0, so we conclude that *η*_3_ is unstable. In the figure, only for the largest λ (top line) is *η*_3_ unstable.

We next analyze stability of the equilibrium points for which *p̅* > 0, labeled *ξ* or 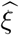 in Figure 2. Stability is equivalent to the trace of *J* being negative and its determinant positive [27, 55], that is:

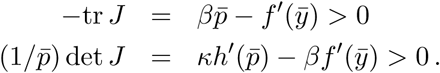

Since *h* is increasing and all constants are positive, these expressions are both positive when *f′*(*y̅*) < 0. Remembering again that the blue curve is an “upside down” version of *f*, we conclude that all the intersections *ξ* shown in Figure 2 are stable, with the possible exception of the intersection labeled 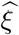, which needs further analysis. At the point 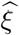, the slope of the green line is *− κ/β* and the slope of the blue line is the derivative of *h*^-1^(−*f* (*y*)) evaluated at *y* = *y̅*, where *h*(*p̅*) + *f* (*y̅*) = 0, so this slope is − *f′*(*y̅*)/*h′(p̅*). Since the slope of the green line is larger (less negative) than the slope of the blue line, we have that *− κ/β > − f′(y̅*)/*h′(p̅*), or equivalently *κh′(p̅) − βf′(y̅*) < 0, so the determinant of *J* is negative, which means that 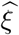 is a saddle point and therefore unstable.

These theoretical results strongly suggest that, for trajectories that start with small *p = u/x*, convergence will be to one of the points labeled *ξ* in Figure 2, or *η*_3_ in the case of the line that does not intersect the blue curve. In other words, if λ, the reproduction rate of the immune challenge, such as a tumor, is small, then we’ll have an intersection below the threshold (dashed line), meaning that the challenge will be eliminated. For a larger, but not too much larger, λ, the intersection is above the threshold, so the challenge will be “tolerated” (it will grow). For λ’ such that there are no intersections with the blue curve, solutions should converge to *η*_3_, which is again under the threshold, implying a second zone of elimination of the challenge. Finally, for very large λ, the immune system is not capable of eradicating the challenge (*ξ* is over the threshold), so escape again occurs. This “two-zone tolerance” (at both intermediate and large λ) is reminiscent of analogous experimental findings, see Box 1.

The analysis based on stability of equilibria can be complemented by numerical simulations, done in Box 1 for an example, as well as a global phase-plane analysis, done below. A simplified case is analyzed further in [59].

### Directions of flow in phase plane

Observe that *ẏ* > 0 in those regions of the phase plane where *h*(*p*) + *f* (*y*) > 0, that is to say, where *p > h^-1^(−f* (*y*)). Geometrically, this means that the flow of the vector field defining the system will point to the right at points that lie over the blue curve (*y* nullcline); it will point left under the blue curve and will be exactly vertical (or an equilibrium) on the curve itself. Similarly, for *p* > 0, *ṗ* > 0 in those regions of the phase plane where *δ_x_* + λ − *κy − βp* > 0, that is to say, where *p < (1/β)(δ_x_* + λ − *κy*). Geometrically, this means that the flow of the vector field defining the system will point up at points that lie under the green lines (*p* nullcline); it will point down over the green lines and will be exactly horizontal (or an equilibrium) on the lines. At *p* = 0, the vector field is horizontal. In summary, motions are “Northeast”, etc., as per these rules in each region delimited by the blue curve and the line *δ_x_* + λ − *κy − βp* = 0:

- NE: over blue, under green
- SE: over blue, over green
- NW: under blue, under green
- SW: under blue, over green

Figure 2 shows the directions of movement in each region, for each of the sample nullclines shown earlier, as well as what a typical trajectory might look like, consistently with these directions of movement and converging to the stable points *ξ* or *η*_3_. (The precise approach depends on the functional forms of *f* and *h* and their parameters. The inset in Figure 2B shows several possibilities.) As λ is increased, the equilibrium shown lies under, above, under, and finally again above the threshold. Next we discuss a specific model where these predictions are verified, and relates our results to experimental evidence of similar phenomena.

## 3 A concrete mathematical model with Treg cells and cytokines

Our results show (1) the role of the underlying IFFL as an estimator of the exponent λ of immune challenge growth and (2) the existence of four regimes, alternating tolerance and rejection, which correspond to different values of λ. In our abstract theoretical arguments, we did not specify the function *f* (*y*), except for the requirement that its graph be as in Figure 1B. We view the model as qualitative and phenomenological, merely as an illustration of surprising behaviors that can arise from, and can be easily explainable by, simple motifs, and not necessarily instantiated by a specific biological system. Nonetheless, it is fair to ask if there is a plausible biological system that gives rise to an *f* (*y*) of this form, and for which our conclusions hold true. We address this issue now, using as a guide simplified versions of standard models in immune dynamics such as found in [38, 34], and using simulations to verify for this system the theoretically predicted four-regime behavior.

We will pick *h*(*p*) = *μp*, a linear function, and 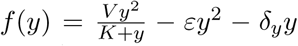. The term 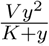 in this expression can be thought of as representing a cytokine-mediated positive feedback, as discussed in Section 7. Motivated by a similar term in [32], the term −*εy*^2^ can be thought of as representing cell-contact-dependent activation-induced cell death (“fratricide”) through the Fas receptor / FasL (“death ligand”) mechanism for T-cell homeostasis suggested by Callard, Stark, and Yates [10]. Finally, the term *−δ_y_y* represents a linear constitutive deactivation and/or degradation. The resulting system is therefore as follows

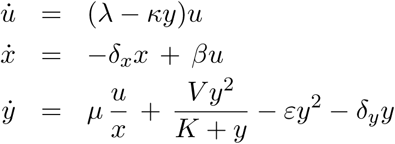

and the reduced system written in (*y,p*) coordinates is

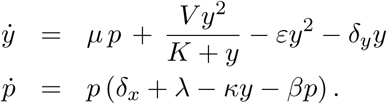

We take the units of time to be days. Cell populations (*u, x, y*) are in units of 10^6^ cells, but *p* = *u/x* is non-dimensional. The parameters that we used, and corresponding units, are as follows: *μ* =10 day^−1^, *V* = 0.25 day^−1^, *δ_x_* = 0.1 day^−1^, *δ_y_* = 0.1 day^−1^, *β* = 1 day^−1^, *ε* = 10^−5^ day^−1^ (10^6^ cells)^−1^, *K* = 100 (10^6^ cells), *κ* = 10^−5^ (10^6^ cells)^−1^ day^−1^ The particular choice of algebraic forms and parameters is discussed in Section 7. Figure 3 shows the nullclines for this system for various increasing values of λ, as well as some typical solution trajectories, showing their convergence to values under, over, under, and finally again over the threshold which determines tolerance or rejection of the immune challenge. This is perfectly consistent with our theoretical predictions.

**Figure 3:**
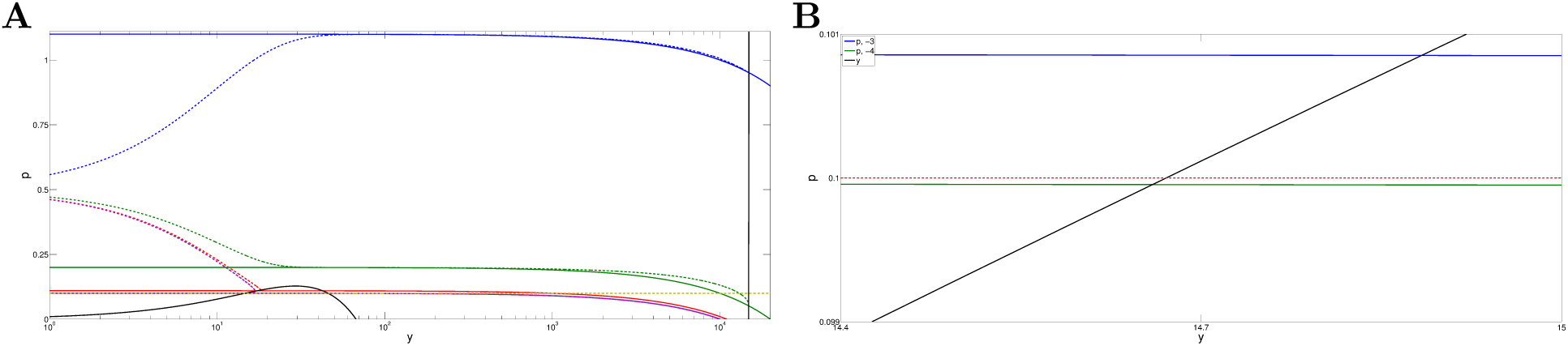
**A.** Phase plane for the system described in the text. Shown are nullclines for several increasing values of λ = 10^*i*^, *i* = −4, −3, −2, −1, 0 (bottom to top). Log scale in *y* is used in order to visualize behavior for different orders of magnitude in λ. Black curve is *y*-nullcline *p* = − (1/*μ*)(*Vy^2^*/ (*K + y*) − εy^2^ − ε_y_y) Solid color curves are *p*-nullclines *p* = (1/*β*)(*d* + λ − *κy*) (which look curved because of log scale), and *p ≡* 0. Horizontal dashed line is threshold *p* = *δ_x_/β* that determines if λ − *κy* is positive or negative. Dashed curves are trajectories, in same color as the respective nullclines. Initial state for simulation is in every case *p*(0) = 0.5, *y*(0) = 0, but only portion of plot for *y* ≥ 10^0^ = 1 is shown. **B.** Zoomed-in view of two nullclines, for λ = 10^-4^ and λ = 10^-3^, to show how steady state falls under/below threshold.

Shown in Figure 4(A-D) are simulations of the complete closed-loop system, where we now included a carrying capacity term for the immune challenge: 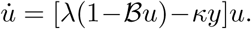 When *u* is small, the term *𝓑u* is dominated by the other terms, but we include this term for numerical convenience in order to avoid blow-up of solutions in the case when *u* is unstable; in any event, such a term is biologically realistic. We picked, specifically, *𝓑* = 10^-3^ (see Section 7). As predicted, for increasing λ there are alternating decay and growth behaviors for *u(t)*. Interestingly, for the third value, λ = 10^−1^, the immune challenge load *u*(*t*) only starts decreasing after about 120 days. This time scale happens to be, purely coincidentally, the start of remission observed in some patients under Ipilimumab [CTLA-4 checkpoint blockade therapy [70]. For this model and parameters, Figure 4(E) shows the asymptotic value of *u*, the immune challenge, as a function of the parameter λ, clearly illustrating the four-regime phenomenon. In Section 8, similar results are shown to hold for a model in which the effect of regulatory variables is through an inactivation of a helper-cell population.

**Figure 4:**
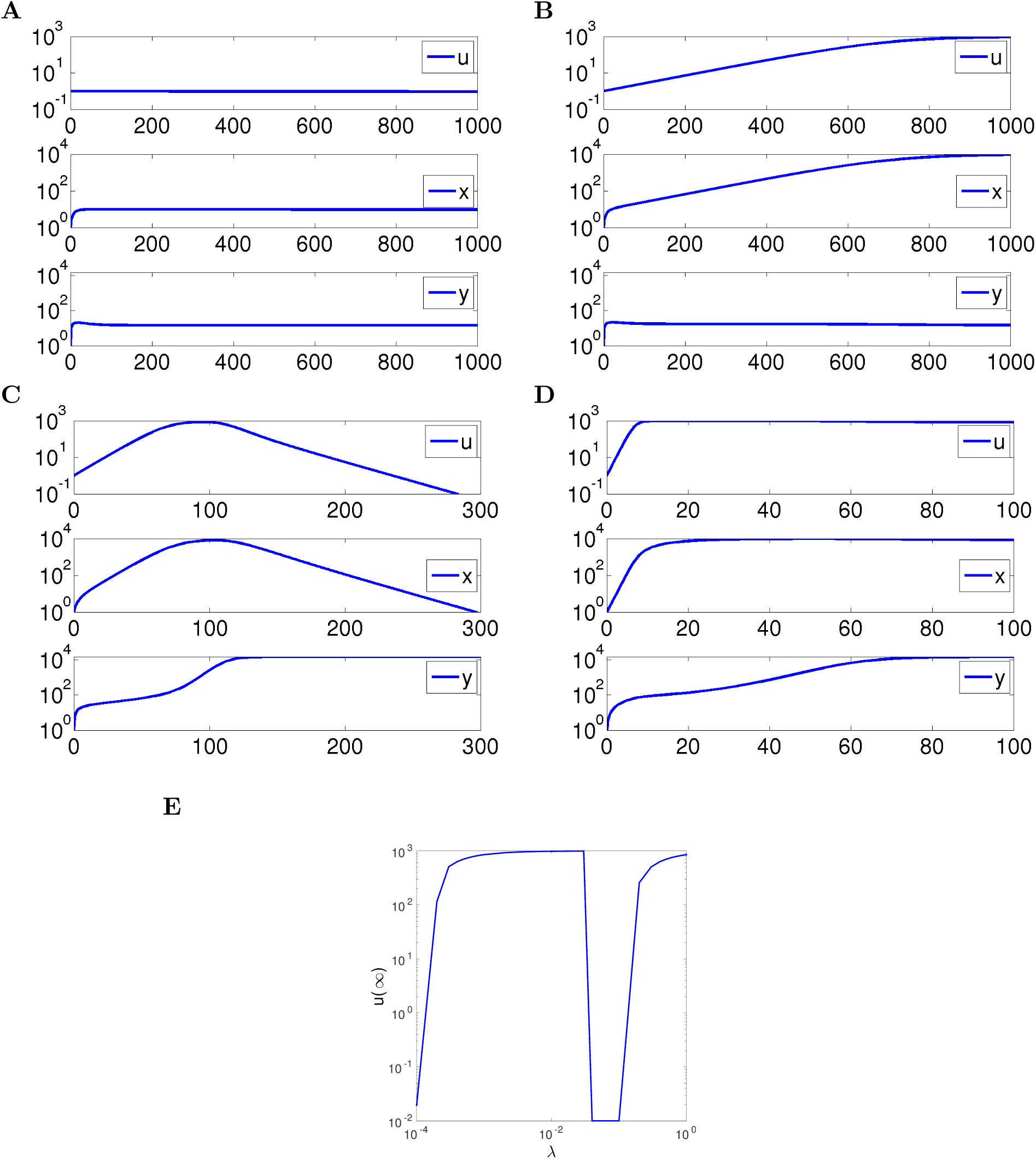
**A-D.** Simulations of full system as described in text, for λ = 10^-4^, 10^-2^, 10^-1^, and 1. (In **A**, *u*(*t*) converges to zero as *t* → ∞, but very slowly.) Parameters as described in text. Initial states are always *u* = 1, *x* = 1, *y* = 0. **E.** Values of *u*(∞) plotted against λ, showing four regimes of elimination, tolerance, elimination, and escape (zero values shown as 10^-2^ to fit in log scale).

## 4 Experimental evidence

We discuss here some experimental findings that appear to be roughly consistent with the behavior of our model.

### Treg cells as a possible regulatory node

Our analysis is merely a phenomenological “toy model” which does not specify immune components. Nonetheless, one might speculate that, as far as T cell activation and deactivation, regulatory T cells (Tregs) may play a role as a regulating intermediate variable *x*. Tregs are a type of CD4^+^ cell that play “an indispensable role in immune homeostasis” [30]. They express surface CD4 and CD25 and internally express the transcription factor FoxP3. Tregs arise during maturation in the thymus from autoreactive cells (“natural Tregs”), or are induced at the site of an immune response in an antigen-dependent manner (“induced Tregs”). They are thought to play a role in limiting cytotoxic T-cell responses to pathogens, and Treg^−^ mice have been shown to suffer from extreme inflammatory reactions. It is known from animal studies that Tregs inhibit the development of autoimmune diseases such as experimentally induced inflammatory bowel disease, experimental allergic encephalitis, and autoimmune diabetes [46]. The various Treg mechanisms can be arranged into four groups centered around four basic modes of action [64]: (1) inhibitory cytokines, including IL-10, IL-35) and TGF-*β*, (2) cytolysis through granzyme-A- and granzyme-B-dependent and perforin-dependent killing mechanisms, (3) metabolic disruption through CD25-dependent cytokine-deprivation-mediated apoptosis, cAMP-mediated inhibition, and adenosinepurinergic adenosine receptor (A2A)-mediated immunosuppression, and (4) targeting dendritic cells through mechanisms that modulate DC maturation and/or function. Moreover, the involvement of T suppressor cells (now Tregs) in regulating the immune response to tumor has a long history, see for instance [20].

### Immune responses to exponentially increasing signals

Our analysis suggests that the immune system might act as an estimator of the rate of growth of an immune challenge, and more specifically the rate of exponential increase. There have been many instances in the experimental literature where related ideas were studied; we mention two of them next.

In their seminal work, Kolsch and coworkers [26] injected exponentially increasing numbers of irradiated syngeneic ADJ-PC-5 plasmacytoma cells into BALB/c mice, starting with 2 cells at day 1, 4 at day two, and doubling subsequent doses for 15 days until about 10^5^ were received. The results pointed to the induction of T suppressor cells (what one now calls Treg cells) as an early event in tumorigenesis that regulated CTL activity. More recently, Kündig and coauthors [29] emphasized the role of antigen kinetics in determining immune reactivity, with exponentially increasing antigenic stimulation recognized by the immune system as a pattern associated with pathogens and thus leading to strong immune responses. They showed that the dynamics of immune stimulation (through dendritic cell vaccination and for T cells stimulated *in vitro*) was by itself a factor in determining the strength of T cell and anti-tumor responses, and “antigenic stimulation increasing exponentially over days was a stronger stimulus for CD8 T cells and antiviral immunity than a single dose or multiple dosing with daily equal doses” concluding that “at a clonal level, T cells are capable of decoding the kinetics of antigen exposure.” They found that IL-2 activation at constant dosage of antigen is almost zero, at linearly increasing dose is higher, and at exponential doses is highest, all roughly consistent with activation of the autocatalytic loop in our model under higher exponential rates, and concluding (Figure 7, caption) that “exponential in vitro stimulation of CD8 T cells enhances IL-2 production and cytotoxicity.” In 2008, Kündig and collaborators, based on this work, obtained a patent [37] for “A method for enhancing T cell response” based on the principle that immunogenicity is enhanced by “exponentially increasing antigenic stimulation of class I MHC CD8+ T cell response… in a manner independent of the dose of the antigen.”

### Rejection of tumors in low and intermediate regions of growth

Another suggestion of our analysis is the existence of intermediate regions of challenge (e.g., tumor) growth in which the challenge will be eliminated by the immune system, with challenges in lower as well as in larger regions not being eliminated. The existence of disjoint regions of tumor elimination depending on rate of growth is very strongly reminiscent of two analogous phenomena, “sneaking through” and “two-zone” tumor tolerance, which have been much discussed since the mid-1960s. The difference is that, in these works, the regions correspond to the magnitude of an initial tumor inocula in animal subjects, rather than growth rates. Nonetheless, there is a surprisingly strong resemblance between our plot in Figure 4 and the ones shown below.

The idea of tumors “sneaking through” from immune control can be traced to the mid 1960s, when Klein [35] found the “preferential take of tumours after small size inocula to a similar degree with that seen with large size inocula, compared to the rejection of medium sized inocula.” Put simply, there is an intermediate region in which tumors can be eliminated. This picture is at least consistent with a larger initial rate of increase in exposure leading to tumor suppression, as in our model. Further, Gatenby, Basten, and Creswick in the early 1980s [21] argued that this four-region phenomenon specifically depends on T-cell repression (just as in our model through the regulatory *x* variable), and framed this role of suppressor T cells on regulating tumor immune response in the more general idea of low zone tolerance (tolerance to antigens under repeated exposure to small antigen doses). To test their ideas, Gatenby et al. carried out experiments that show sneaking through behavior as well as the failure of this behavior when “suppressor T cells” are eliminated, see Figure 4A and Figure 4B respectively. Murine sarcoma Meth A was administered in varying doses to BALB/c mice, and incidence of tumors was measured in each group of 12-42 mice, at two weeks after the last mouse died from tumor. Similar results on sneaking-through had been reported by Kolsch and Mengersen in previous work in which mastocytoma BM3 injected cells were injected into BALB/c mice, see Figure 4C. Care must be taken in interpreting these experimental numbers in terms of a model. The numbers reported are for “tumor incidence,” meaning percentages of mice in which tumors were detected by some predetermined point. If we assume that survival (until mouse sacrifice, or indirect death from the tumor) depends probabilistically on tumor size, then we could think of tumor incidence as a proxy for size.

McBride and Howie [43] also found experimentally what they and several other authors refer to as “two-zone tolerance,” meaning the existence of two separate regions (low and medium ranges) where a tumor can be eliminated, and, again as in Gatenby et al.’s work, they focused on the role of suppressor cells in the phenomenon. The authors argue that their results “support the view that immunogenic tumours, as they grow from small cell numbers, might be able to escape host surveillance by specifically tolerizing the immune system… [and] large tumour burdens can interfere with the host’s immune response by inducing suppressor cells.” The authors studied the effect of injecting fibrosarcoma (FSA) cells in C3Hf mice at various numbers, and then quantified immune effect on tumors by extracting splenic T cells, mixing them with additional viable FSA, and injecting the mix into new mice, finally measuring after a certain predetermined time the number of sites in target mice where tumor growth was detected (out of 40 sites in 20 mice).

Both [21] and [43] used in their protocols exponentially increasing tumor doses for different animals, but neither used, apparently, exponential dose escalation in time, notwithstanding the statement in [43] that their work “confirms and extends the findings of Kolsch and coworkers… when irradiated plasmacytoma cells are injected in… exponentially increasing doses.” (Indeed, as we already discussed, [26] used exponential dose escalation in time.) Since there are many unmodeled processes involved in antigen presentation and processing, the injection of a certain number of cells can be viewed as providing an effective input time-varying signal whose slope depends on this exponentially varying injection amount. Nonetheless, experiments to test the explicit dependence on exponential rates remain to be performed. See also our alternative model in Section 9.

We also remark that somewhat analogous experiments have been performed for interactions of the immune system with viruses instead of tumors. Notably, Bocharov and colleagues [6] studied the effect of varying exponential rates of growth, λ in our notations, of Hepatitis B and Hepatitis C viral infections, and found non-monotonic responses in immune responses and clearance rates.

## 5 Comparison to other models

We next briefly discuss our interpretation of some existing models for self/nonself dynamic discrimination in immune systems, and compare them to the IFFL model.

### Tunable Activation Threshold (TAT)

This model was suggested by Grossman and Paul [24] in 1992, motivated by the realization that “self/nonself discrimination may be much more complex than the simple failure of competent lymphocytes to recognize self-antigens”. The authors argued that for a stimulus to cause cell activation, the excitation level must exceed an activation threshold, and when engaged in persistent sub-threshold interactions, cells are protected against chance activation. In the TAT model, an *activation threshold* for an immune cell is dynamically modulated by an environment-dependent recent excitation history. This history is summarized by an *excitation index,* which we will denote as *x*(*t*), which computes a sort of weighted average of the cell’s past excitation levels. Given temporal excitation events, which we denote by *u(t),* it is assumed that the cell undergoes perturbations that depend on the difference between *u*(*t*) and the memory variable *x*(*t*). The key assumption is that such a perturbation, which we write as *y(t)*: = *u*(*t*) − *x(t)*, must exceed a fixed critical value, which we denote by *θ*, in order to cause activation. In other words, it must be the case that *u*(*t*) − *x*(*t*) > *θ*, or equivalently, *u*(*t*) > *x(t)* + *θ* (this is how we interpret the statement in [24] that “the activation threshold equals the excitation index plus that critical value”) for activation to occur. Cells maintained at a high level of excitation *x*(*t*) therefore are relatively insensitive to activation, thus being in some sense anergic. The authors deduce from their model that “upon gradual increase in the levels of excitation…a cell is not likely to be activated…it will become progressively anergic” which is intuitively equivalent to our remark about the lack of continued excitation under slow increases in antigen presentation. With our notations, the model suggested in [24] is:

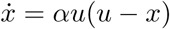

for some constant *α*, and the output would be *y = u − x*. (No explicit population-based nor signaling mechanism was given.) Notice that we then can derive a differential equation for *y*:

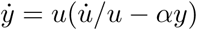

and this means, roughly, that *y* should approach 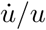, the logarithmic derivative of the input *u*, so that a “log sensing” property is satisfied by the output. Moreover, when the input is constant, the output converges to the same value (zero), independently of the actual value of the input, so we have perfect adaptation. Moreover, we expect *y* to be small (and thus not exceeding the threshold θ) unless *u* changes fast in the sense that its logarithmic derivative is large. For example, for *u*(*t*) increasing linearly, 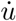 would be a constant, so 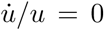 and therefore *y*(*t*) → 0 as *t* → ∞. On the other hand, for an exponentially increasing *u*(*t*), *y*(*t*) converges to a value proportional to the exponential rate. These properties are analogous to those satisfied by our model.

### Discontinuity theory of immunity

This model was suggested by Pradeu, Jaeger, and Vivier [48] in 2013 as a “unifying theory of immunity”. Their key hypothesis is that effector immune responses are induced by an “antigenic discontinuity” by which they mean a “sudden” modification of molecular motifs with which immune cells interact. The authors present evidence that natural killer (NK) cells and macrophages are activated by transient modifications, but adapt (ceasing to be responsive) to long-lasting modifications in their environment, and then propose to extend this principle to other components of the immune system, such as B cells and T cells. They also argue that although tumors give rise to effector immune responses, “a persistent tumour antigen diminishes the efficacy of the antitumor response”. In summary, their criterion of immunogenicity is the phenomenological antigenic discontinuity and not the nature of the antigen, including both “discontinuities” arising from self motifs such as tumors as well as from non-self motifs such as bacterial or viral infections. As examples of mechanisms for desensitization they mention receptor internalization, degradation or inactivation of signaling proteins. A concrete example of the latter is the dephosphorylation triggered by immunoreceptor tyrosine-based inhibition motif (ITIM)-containing receptors antagonizing kinases triggered by immunoreceptor tyrosine-based activation motif (ITAM)-containing receptors. The authors also mention Treg population dynamics.

Using our notations, the model in [48] starts by computing a running average of the absolute value of consecutive differences in inputs presented at discrete times on a sliding window *K* time units long:

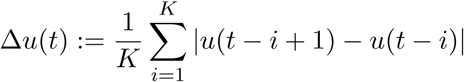

and then taking as the output *y* = Θ(Δ*u*), where Θ is a sigmoidal saturating function. The authors employ

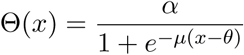

but one could equally well (and perhaps easier to justify mechanistically) employ a Hill-type function 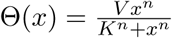. In continuous time, and assuming that the input is differentiable, we could interpret

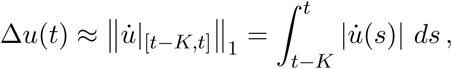

where the right hand term is the total variation of the input on this sliding window. Note the absolute value, which means that, in this model, activation is symmetrically dependent on increases or decreases of the excitation: decreases may help with “missing self” recognition, in which the expression of a “self” marker suddenly decreases, thus triggering a response. As with our model, slow variations in the input will lead to small *y*(*t*), with the threshold function Θ resulting in an ultrasensitive, almost binary, response (provided that μ or n are large, in the two suggested functions Θ).

### Growth threshold conjecture

This model was suggested by Arias, Herrero, Cuesta, Acosta and Fernández-Arias [4] in 2015 as “a theoretical framework for understanding T-cell tolerance” based on the hypothesis that “T cells tolerate cells whose proliferation rates remain below a permitted threshold”. As in the other works, the authors postulate that T cells tolerate cognate antigens (irrespectively of their pathogenicity) as long as their rate of production is low enough, while those antigens that are associated with pathogenic toxins or structural proteins of either infectious agents or aggressive tumor cells are highly proliferative, and therefore will be targeted as foes by T cells. In summary, once again the postulate is that a strong immune response will be mounted against fast-growing populations while slow-growing ones will be tolerated. The model in [4] is not one of change detection as such, but it is a closed-loop system that includes both detection and a killing effect on pathogens. To compare with our previous models, let us again denote the pathogen population size (or a density in a particular environment) by *u*(*t*) and the effector cell population by *y*(*t*). The authors give for *y* a second order equation *ÿ = −δy + αu*, modeled on a spring-mass system that balances a “restoring to equilibrium force” to its activation by pathogens. We prefer to write the system as a set of first order ODE’s. Thus, we let *x: = ẏ*, and write:

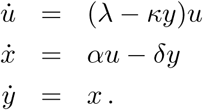

The *u* equation has an exponential growth term balanced by a kill rate that depends on the effector population. The effector population integrates the amount of *x* (which we might interpret as an intermediate type of cell); the growth of *x* is driven by pathogens, with a negative feedback from *y* (in essence an integral feedback on *x*), but there is no obvious biological mechanism for this model. Observe that when there is no pathogen, this results in a harmonic oscillator for *x* and *y*, with sustained oscillations and even negative values. In any event, the authors computationally obtain a bifurcation-like diagram in the (λ, *κ*) plane, dividing this plane into two regions, labeled “tolerance” (of infection, hence, failure of the immune system) and “intolerance”. These regions show how to trade off the growth rate λ of the pathogen versus the parameter κ, which represents a combination of affinity and clearance rate, and various conclusions regarding evasion strategies and the role of fever and even Treg cells are qualitatively derived from there.

## 6 Perfect adaptation and scale-invariance

A system is said to be *perfectly adapting* provided that its response returns asymptotically to a pre-stimulus value under constant stimulation. This property is typically exhibited by sensory systems processing light, chemical, and other signals, and it has been extensively investigated both experimentally and mathematically [2, 31]. In particular, when subjecting a perfectly adapting system to a step-wise input signal, as shown in Figure 6, the output of the system settles, after a transient response, to a basal value which does not depend on the magnitude of the stimulus. The response amplitude and timing, on the other hand, typically depends on the input magnitude. This notion can be refined as follows. Suppose that every step has the same relative or “fold” change, *u_i+1_ /u_i_* = constant, as shown in the figure. For *scale-invariant* systems, the responses to such steps have the exact same shape, amplitude, and duration.

**Figure 5:**
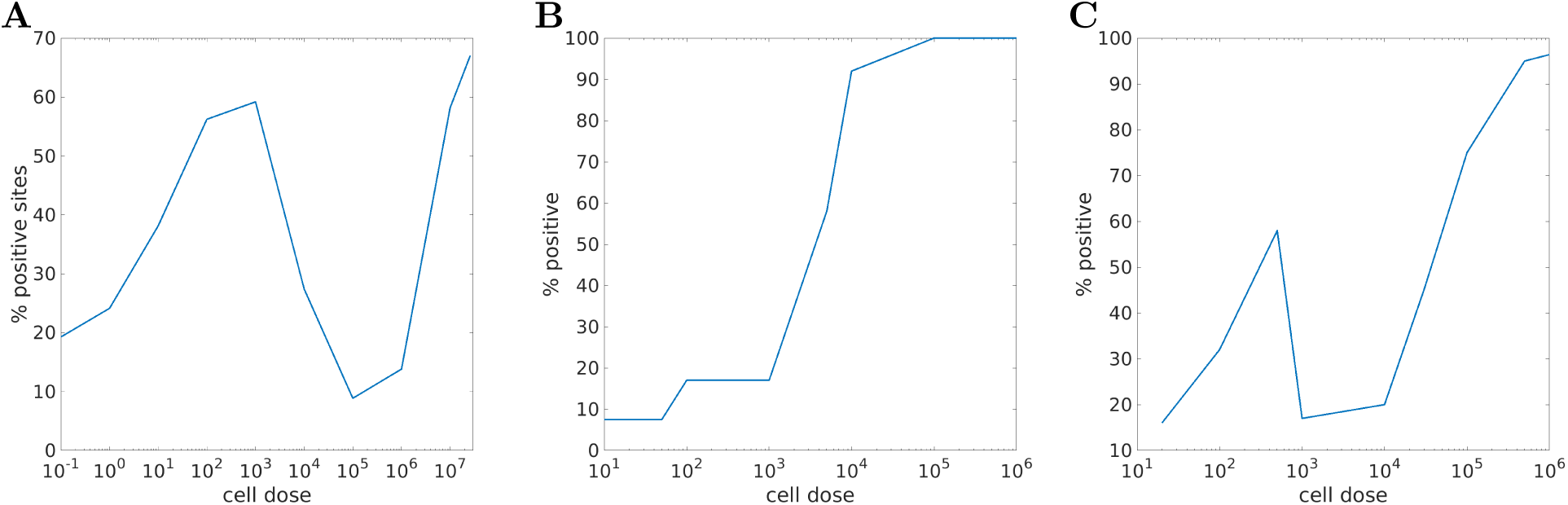
**A**: four regions of “sneaking through”, and **B**: no sneaking-through when eliminating suppressor T cells, plots drawn from data in [21]. Very low zone (less than 100 cells: low tumor incidence; low zone (10^2^—10^3^ cells): tolerance of tumor; moderate zone (10^3^—105 cells): immunogenic (low tumor); high zone (>10^6^): again tolerogenic. **C**: Another illustration of the same general idea of four regions. Plot drawn from data in [36]. The *x* axis in these plots case measures the initial number of tumor cells, and not the rate of growth λ of tumor as it did as in Figure 4. See text for details on tumor challenge and response criteria.

**Figure 6:**
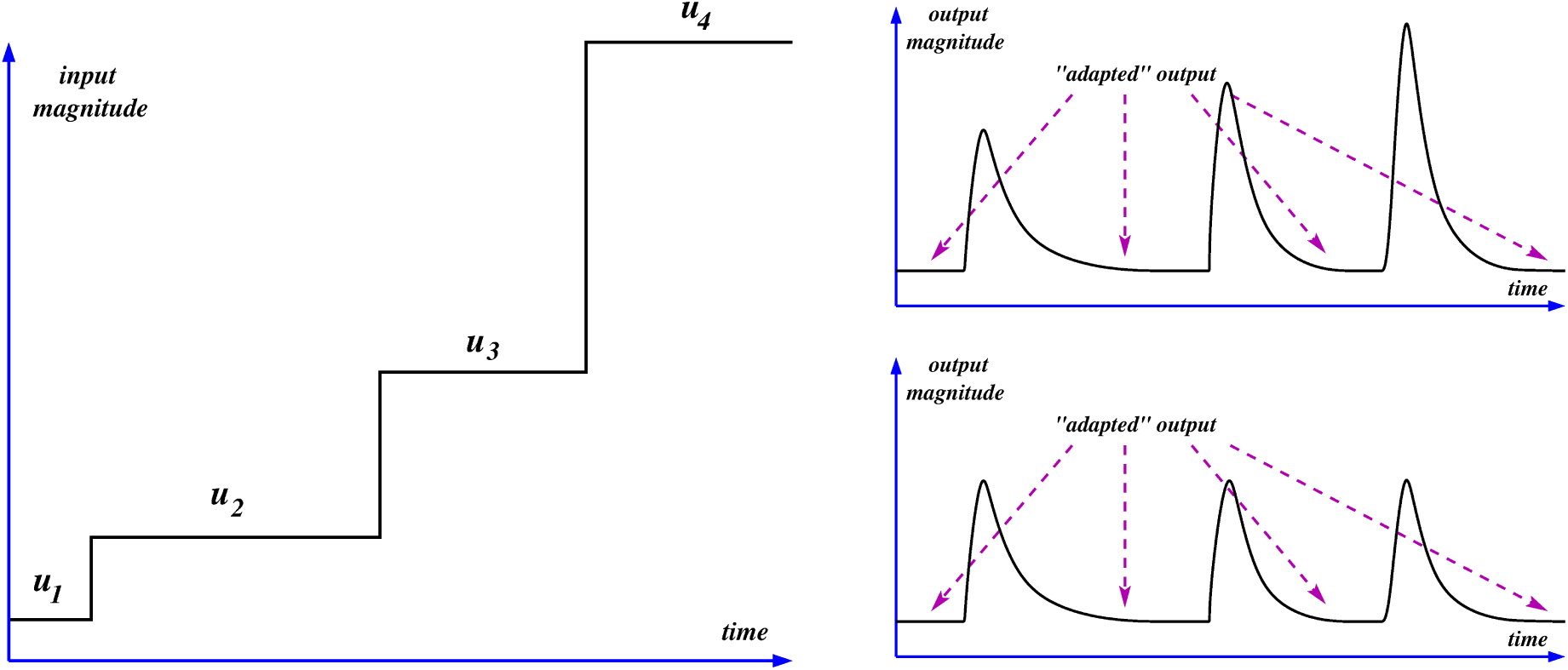
In a perfectly adapting system, a step-wise input (left) gives rise to different responses that settle to the same basal level (top right). If the system has the scale invariance property, these responses are identical (bottom right)

The alternative term “fold change detection” is sometimes used for this property, to emphasize the fact that such systems can only react differently if the fold changes are not the same. To put it in another way, such systems can give different responses if difference log *u_i+1_* −log *u_i_* is nonzero (log sensing) as opposed to *u_i+1_ − u*_i_. The precise mathematical definition of scale-invariance involves arbitrary input signals: responses to arbitrary scaled inputs as in Figure 7, and not only piecewise constant ones, should be the same, provided that the internal state starts from a preadapted value. We refer the reader to [51] for technical details.

**Figure 7:**
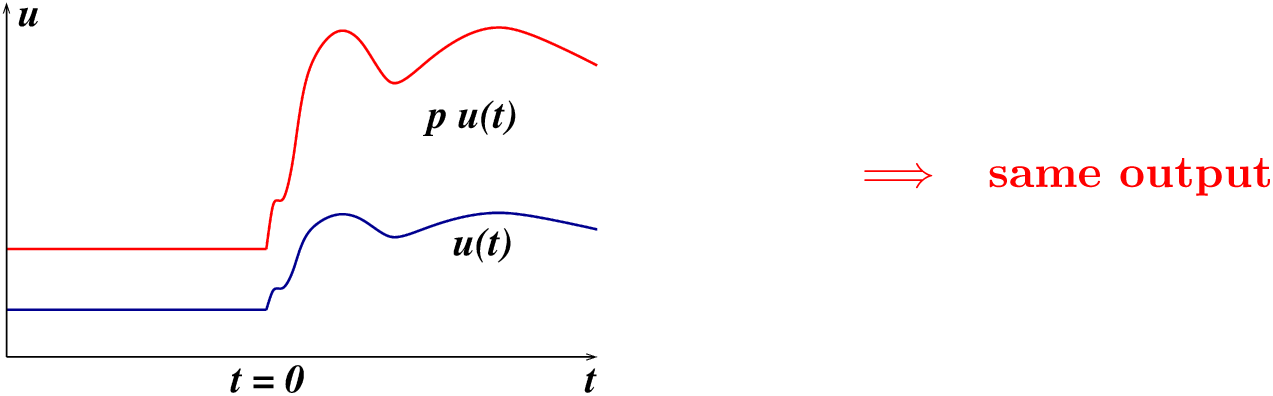
Scale invariance means that scaled signals should result in the same output, provided that the initial state is preadapted to the respective constant value for *t* < 0

Scale invariance or fold change detection (FCD) is a strengthening of the Weber-Fechner “log sensing” property, which is sometimes defined as the requirement that the maximum amplitude of responses to two scaled inputs should be the same, but not necessarily their exact shape or even timing. Recent interest in the FCD property was largely triggered by the papers [22, 12], in which fold-change detection behavior was experimentally observed in a Wnt signaling pathway and an EGF pathway, respectively; these are highly conserved eukaryotic signaling pathways that play roles in embryonic patterning, stem cell homeostasis, cell division, and other central processes. Later, the paper [52] predicted scale invariant behavior in *E. coli* chemotaxis, a prediction which was subsequently experimentally verified [40]. Similar results are available for other bacterial species, for example *R. sphaeroides,* for which theoretical predictions made in [25] were experimentally confirmed in [66]. A mathematical study of scale invariance, together with a necessary and sufficient characterization in terms of solutions of a partial differential equation, can be found in [51]. It has been recently shown that all scale invariant systems compute a certain type of differentiation operator, such as logarithmic derivatives [39].

One example of a scale invariant system is the IFFL (2) that underlies our model, which we repeat here for ease of reference:

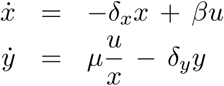

where *β, μ, δ_x_, δ_y_*, are some positive constants and *u*(*t*) is viewed as an external stimulus. For any given input function *u*(*t*) and initial values *x*(0) and *y*(0), the solution of this system can be found by first solving the scalar linear ordinary differential equation for *x*(*t*), and then plugging this result together with *u*(*t*) into the *y* equation, which is also a linear ODE. For a constant input *u*(*t*) ≡ *u*_0_ > 0, there is a globally asymptotically stable steady state, given by

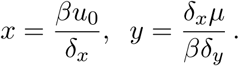

At steady state, the output *y* is independent of the particular value of the constant input *u*_0_, meaning that the system is perfectly adapting. Suppose next that (*x*(*t*), *y* (*t*)) is any solution of the system (2) corresponding to an input *u*(*t*), now not necessarily a step function. It is then immediate to verify that (*px*(*t*),*y*(*t*)) is a solution corresponding to the input *pu*(*t*), *t* ≥ 0, for any nonzero constant scaling factor *p*:

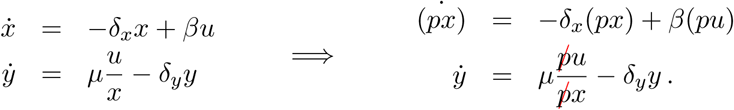

Thus, this system responds with the same output signal *y*(*t*) to two inputs which differ only in scale, provided that the initial state *x*(*t*) had already adapted to the input at time *t* < 0. In other words, given a step input that jumps from *u*(*t*) = *u*_0_ for *t* < 0 at time *t* = 0 and an initial state at time *t* = 0 that has been pre-adapted to the input *u*(*t*) for *t* < 0, *x*(0) = *βu*_0_/*δ_x_*, the solution is the same as if, instead, the input would have been *pu*(*t*) for *t* > 0, but starting from the respective pre-adapted state *pβu*_0_/*δ*_*x*_. This means that our IFFL subsystem is scale-invariant.

It would be very interesting to test experimentally the response to scaled versions of antigen presentation, to verify if such scale invariance holds, even in an approximate fashion.

## 7 Details on the model used for simulations

In this section, we explain the terms in the differential equations used in simulations, including the parameters used. Of course, our model is only a cartoon of a hugely complicated system of interlocking processes. Moreover, even if the model were mechanistic, which it is not, numbers would depend on the specific tumor or infection tissue being modeled. Thus, these algebraic forms and numbers are offered only as a plausible scenario.

As explained in the main text, *u* represents an immune challenge, specifically a tumor in this case, while *x* and *y* might represent populations of activated and specific T suppressor (CD4^+^ CD25^+^ Treg) and cytotoxic T cells (CD8^+^ cells) respectively. We use as a guide in our modeling the paper by Kirschner and Panetta [34], which has become a classic reference for tumor-immune interactions in the presence of cytokines (no regulatory T cells in that model), together with the more recent paper by Khailaie et al. [32] which described a model of immune activation in the presence of both chemokines and also regulatory T cells (no tumor dynamics in that model).

### Cell number units

Since we use parameters from both [34] and [32], it is thus important to clarify the units used in these sources.

Kirschner and Panetta’s paper gives “volume” as the unit for cell populations. Since many of these parameters were in turn obtained from the foundational paper by Kuznetsov et al. [38], which provided one of the first differential equation models for interactions between tumors and the immune system, one can compare the two papers, to map their unit to cell numbers. For this purpose, we can compare the value of the carrying capacity of tumors (***“𝓑”*** in the simulations that we provided) in both papers. In [34] ***𝓑*** = 10^−9^, and in [38] ***𝓑*** = 2 × 10^−9^. Ignoring the factor of 2, this means that “volume” = number of cells. This is confirmed by comparing the Michaelis-Menten constant for IL-2 activation *g*1 (*g* in the second paper): 2 × 10^−7^ volume units and 2.019 × 10^−7^ T cells respectively. Therefore, we will be interpreting cell units in [34] as numbers of cells. In our simulations, we use ***𝓑*** = 10^−3^, because we prefer to switch to units of 10^6^ cells.

In Khailaie et al.’s paper (personal communication from first author), “cell” means nondimensional units, cells/*C*_0_, where *C*_0_ is an unspecified reference quantity of cells. Now, Figure 5 in [32] shows stable branches of equilibria under antigen stimulation in ranges of 2 to 30 nondimensionalized T cells, while in their companion experimental paper [44], the same authors provide estimates of T cells in various tissues in mice in the range 10^6^ to 8 × 10^6^. Thus approximately *C*_0_ = 10^6^ cells is consistent with the analysis in [32], and so we will interpret the numbers in that reference in units of 10^6^ cells.

### The autocatalytic term 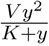

This term is intended to model a cytokine-mediated positive feedback loop on effector T cells. Cytokines are molecules that act as immunomodulating agents and mediate communication among immune systems components and their environment. Their concentrations can increase up to 1,000-fold during inflammatory conditions. Examples of cytokines include interleukins such as IL-2 and IL-6, interferons, and TNF. The role of cytokines in anti-tumor responses, and in particular IL-2, has been the subject of much study [14] and of mathematical modeling since at least the work of Kirschner and Panetta [34], who proposed a simple differential equation model that includes variables for tumor load, effector immune cells, and cytokines. In their model, activated T cells produce cytokines, specifically IL-2, which in turn enhance lymphocyte activation, growth and differentiation, in particular of the cytotoxic T cell (CTL) population. The effect is through a positive feedback that is both autocrine, that is, acting on the cells that produce it, and paracrine, acting on nearby cells. This role of IL-2 in enhancing T-cell proliferation and differentiation is one reason that IL-2 was originally named “T-cell growth factor,” although by now many other immunoregulatory functions of IL-2 are known.

The term that represents the effect of the cytokine (IL-2) on *y* in [34] is *p*_1_ *yz*/(*g*_1_ + *z*), where the cytokine *z* satisfies the differential equation *ż* = *p*_2_*uy*/(*g*_3_ + *u*) − *μ*_3_*z*. This equation models IL-2 secretion by activated effector T cells, with a Michaelis-Menten kinetics to account for self-limiting production of IL-2, together with a degradation rate. To obtain z as a function of *y*, we assume that this variable is at equilibrium; on the saturation regime of antigen load *u* we obtain *z* = (*p*_2_/*μ*_3_)*y*. Now substituting this expression into the differential equation for *y*, we have the autocatalytic term

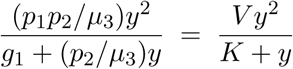

where *V* = *p*_1_ and *K* = *μ*_3_*g*_1_/*p*_2_ If we start, instead, from [32], the corresponding term in the differential equation for *ẏ* is *ayz*, where *z* now satisfies a different equation, *z′* = *dy − eyz − fz* and the term *eyz* represents IL-2 consumption rate by T cells. Nonetheless, under the same equilibrium assumptions we obtain *z = dy/(f + ey)*, which when substituted into *ayz* gives

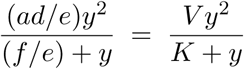

where *V = ad/e* and *K = f/e*. In other words, we derived the same functional form as when starting from [34].

Plausible parameters values can be obtained from [34] or from [32]. The parameters used in [34] were *p*_1_ = 0.1245, *p*_2_ = 5, *g*_1_ = 2 × 10^7^, and *μ*_3_ was arbitrarily picked as 10 from the range 8.31 to 33.27 using a half-life for IL-2 of 30 to 120 minutes given in [50]. Plugging these into the formulas given above, we obtain *V* = 0.1245 and *K* = 10^6^*K*_0_, where *K*_0_ ranges from 33 to 133. As discussed earlier, we are reading the units in the paper [34] as individual cell counts. When translating to our units of 10^6^ cells, we obtain that *K* in their model ranges between 33 and 133. (The argument is: if we rescale variables letting *η* = *y*/10^6^, then the corresponding term in 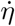 is 10^−6^*V*(10^6^*η*)^2^/(10^6^*K*_0_ + 10^6^*η*) = *Vη*^2^/(*K*_0_ + *η*), which means that *K = K_0_* when writing the equation in terms of *η*.) Using parameters from [32] gives similar results. As discussed earlier, we are reading the units in that paper as 10^6^ cells. These parameters are picked in [32] as follows: *a* = 0.4, *d* = 0.01, *e* = 0.01, *f* = 1. Plugging these into the formulas given above, these lead to *V* = 0.4 and *K* = 100. In summary, one paper gives *K* between 33 and 133 and *V* = 0.1245, and the other paper uses *K* = 100 and *V* = 0.4. We therefore take *K* = 100 and for *V* pick an average, *V* = 0.25 of the two values. Note that the units of *K* are 10^6^ cells, and the units of *V* are day^-1^.

### The fratricide term *−εy^2^*

Following the T cell model in [32], we include the term *−εy^2^* for cell-contact-dependent activation-induced cell death in activated T cells, a process known as “fratricide”. Activated T cells express the receptor FasR, also known as apoptosis antigen 1 (APO-1 or APT), cluster of differentiation 95 (CD95) or tumor necrosis factor receptor superfamily member 6 (TNFRSF6), as well as the ligand for this molecule, FasL; fratricide can result from direct cell contact or from cleavage of FasL (“death ligand”), and the ligation of FasR by soluble FasL results in apoptotic cell death, mediated by caspase activation [19]. It is believed that the exposure to tumor antigens in T cells might mediate fratricide [41]. Callard, Stark, and Yates [10] modeled the fratricide mechanism by a nonlinear death term *−εy^2^* and speculate that Fas-mediated apoptosis results in a density-dependent death rate for T cell homeostasis that does not require competition for resources nor quorum-sensing mechanisms for density estimation. From [32], we pick *ε* = 10^−5^, in units of day^−1^ (10^6^ cells)^−1^.

### The decay terms *− δ_x_x* and *δ_y_y*

These represent linear degradation of activated T and Treg cells. The values *δ_x_x* = *δ_y_y* = 0.1 are from [32]. Units of both are day^−1^.

### The term *βu*

Stimulation of regulatory cells is a very complex process that involves a wide variety of antigen presenting cells and other mediators. TRegs are exported from the thymus and recirculate through secondary lymphoid tissues as “central” TReg cells, and get activated through T cell receptor (TCR) ligation, CD28 co-stimulation and/or interleukin-2 (IL-2), which induce upregulation of expression of interferon regulatory factor 4 (IRF4), which then orchestrates their differentiation into “effector” TReg cells [42]. We make the simplest possible assumption: the rate of activation is proportional to the immune challenge such as a tumor population, that is, we postulate a term *βu* in *ẋ*. It is virtually impossible to give a numerical value for the parameter *β*, since this value depends on the nature of the immune challenge, spatial relations between antigen presenting cells and T cells, and so forth. Khailaie et al. [32] simply use a term *+k(t)* to represent this stimulation (where *κ(t)* is the product of antigen stimulation “*β*” and the supply *N* of naive T cells or resting Treg cells, and introducing an unspecified multiplier to model possibly different effects on T cells compared to Tregs). This additive input is naturally modeled by *β = 1*, and we take that simplest possible value. Units are day^−1^.

### The term *μu/x*

There are various ways to justify this term. We picked a mathematical form for the effect of the immune challenge *u* and regulatory elements *x* on effector cells *y* that is the simplest possible to model activation by *u* and repression by *x*. Let us discuss why this choice is reasonable phenomenologically. The term “regulatory T cell” (Treg) actually encompasses several subclasses of cells that help in peripheral tolerance, preventing autoimmune diseases, and down-modulating immune responses. These cells they affect many other immune components, from B cells to helper cells (Th1, Th2, Th17) and cytotoxic T cells, through both direct and indirect interactions. These interactions form an extremely complicated and poorly understood network that includes inhibitory molecules such as CTLA4 and messaging by cytokines (TGF-*β*, IL-10, IL-35, and others) which result in the suppression of helper cell differentiation and in indirect down-regulation of MHC and costimulatory molecules on antigen-presenting cells, thereby reducing T cell activation. The repression of T cell activation through TCR-MHC is one way to see the negative effect of *x* on *y*. Another is the indirect effect through inhibitory cytokines such as IL-10, TGF-*β*, and IL-35 that can suppress T cell activation. The simplest mass-action kinetics model would assume independent effects: activation by *u* and repression by *x*, leading to a term of the form *h*_1_(*u*)*h*_2_(*x*) driving *y* activation. For the effect of *u*, let us take *h*_1_(*u*) = *c*_1_ *u*, for some constant *c*_1_. If we assume that *x* cells (or messenger molecules) repress through binding to a certain type of receptor, and *R* represents the number (or fraction, or concentration, depending on units) of free receptors, then at equilibrium we would have *κRx = R*_0_, where *R*_0_ quantifies occupied receptors, and from a conservation *R* + *R*_0_ = *R_T_* assuming a constant total number of receptors, we would have that *R = R_T_/(1 + kx)* is the number of free (unbound) receptors, so unless *κ* << 1 we may take *h*_2_(*x*) = *c*_2_/*x* for some constant *c*_2_. These arguments will result in the algebraic form *h(x, u)* = *Mu/x*.

A different justification is as follows. Let us assume that there is an intermediate variable *z*, which might represent for example a population of helper T cells (Th cells or CD4+ T cells) which helps activate the cytotoxic T population *y* and is itself activated by the immune challenge *u* and inactivated by the regulatory variable *x*. The simplest equation would be *ż = −μ_0_xz+βu* where we are assuming that helper cells are also being activated in a manner proportional to the magnitude of the immune challenge, and *μ_0_x* represents the *x*-dependent degradation of *z*. We assume that *ẏ* has a term *z* corresponding to activation by helper cells. Assuming that this equation is at equilibrium, we may substitute *z = (*β*/μ_0_)u/x* into the *ẏ* equation, giving a term *μu/x*, where *μ* = *β*/*μ*_0_. (If helper and T cell activations are at similar timescales and the equilibrium assumption is not made, one add may the *z* differential equation explicitly. We prefer to keep the model simpler, but see Section 8 for simulations using that model.) Khailaie et al. [32], include in T cell dynamics a similar mass-action degradation or inactivation term, using a rate constant 0.1. Following this, we pick the value *μ*_0_ = 0.1, so that, together with *β* = 1 we have *μ* = 10. As *u* and *x* are both in units of 10^6^ cells, *μ* has units day^−1^.

### The terms *λ_u_* and *−κyu*

The term *λu* is a standard exponential growth term. We view λ as a varying parameter, which quantifies the initial exponential growth of the immune challenge.

The killing term *−κuy* in the 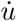 equation represents a simple mass-action suppression of the immune challenge, such as cytotoxic T cells killing tumor cells. The constant *κ* depends on many factors, such as the type of tumor, size and geometry of tumor microenvironment, accessibility of tumor cells to vasculature, and so forth. In the original paper by Kuznetsov et al. [38], one finds *κ* = 1.101 × 10^−7^ in units of day^−1^ cells^−1^ which when normalized to units of 10^6^ cells would give the value *κ* = 1.101 × 10^−1^, This value seems to be too large for most cancers. For example, based on fits to experimental data, the recent paper [67] obtains a number which is many orders of magnitude smaller. That paper analyzes the killing by cytotoxic CD8+ T cells of MHCI^+^ tumor cells in a B16 mouse metastatic melanoma model, and determines a killing term for such cells of the following form (with different notations here): *− [c/(ε + U)]Yu*, where *Y* is the concentration of effector CD8^+^ T cells in the tumor microenvironment, using units of cells/mm^3^, *c* is a constant that quantifies MHCI positive tumor death rate due to T effectors, and has the value 2.49 × 10^−13^ in units mm^3^ day^−1^, *U* is the total number of tumor cells, *ε* is a “small number” to account for other cells, and *u* is the number of major histocompatibility complex class I positive tumor cells. Since [*c/(ε + U)]Y* has units day^−1^, if we convert to *y* in units of 10^6^ cells, we obtain *cY = κy* where *κ* = 2.49 × 10^−7^/(ε + *U*) has units (10^6^ cells)^−1^ day^−1^. Depending on the number of cells *U* in the tumor, this number *κ* could be very small, and it is certainly less than 2.49 × 10^−7^. To take another example, Kirschner and Panetta [34] employ a Michaelis-Menten killing term *−auy/(g_2_ + y*), with *a* = 1 and *g*_2_ = 10^5^. Given these wide ranges, we pick *κ* = 10^-5^ for our simulations. Units are (10^6^ cells)^−1^ day^−1^. (A two-zone behavior of tumor elimination can also be found with *κ* = 10^−4^, *κ* = 10^−3^, *κ* = 10^−2^, and *κ* = 10^−1^, but shifting the range of λ’ at which different behaviors arise.)

### Sensitivity to parameters in the function *f*

We recall the definition of the function *f*:

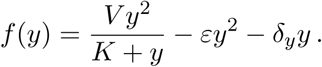

The main requirement for the theoretical analysis in the main text is that *f* have a cubic form as illustrated in Figure 1, so that then the nullcline analysis in Figure 2 applies. In other words, *f* should have one zero at *η*_1_ = 0 and two positive zeros *η*_2_, *η*_3_ so that *f(y)*< 0 for *η*_1_ < y < *η*_2_, *f* (*y*) > 0 for *η_2_ < y < *η*_3_*, and *f* (*y*) < 0 for *η*_3_ < *y*. (Observe that signs gets reversed in the nullclines in Figure 2, because of the negative sign in the formula *p = h^-1^(−f(y)*).) Writing 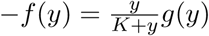 where

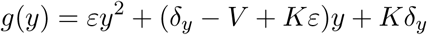

and using that 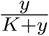 is positive for *y* > 0 and zero at *y* = 0, the requirements on *f* translate into the requirement that the parabola *g*(*y*) have two positive zeros *η*_2_, *η*_3_ (and be negative in between them), which is equivalent to:

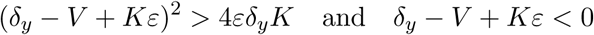

For our parameters, *V* = 0.25, *K* = 100, *δ_y_* = 0.1, *ε* = 10^-5^, we have *δ_y_* − *V* + *Kε* ≈ −0.1490, (*δ_y_*− *V* + *K*ε)^2^ ≈ 0.0222, and 4*εδ_y_K =* 4 × 10^−4^, so that these conditions are satisfied. These requirements imply that the maximal autocatalytic strength *V* should be large, and the degradation constant *δ_y_* and the fratricide constant *ε* should be small.

## 8 A model with an intermediate population

We consider here that a slightly different model, in which *u* and *x* affect the effector variable *y* only indirectly, through production and repression respectively of a “helper cell” population. Figure 8 plots simulation results (all parameters exactly the same as in earlier model), showing that this model leads to similar results as those for the simpler model.

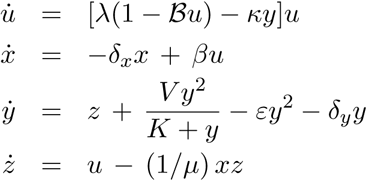

**Figure 8:**
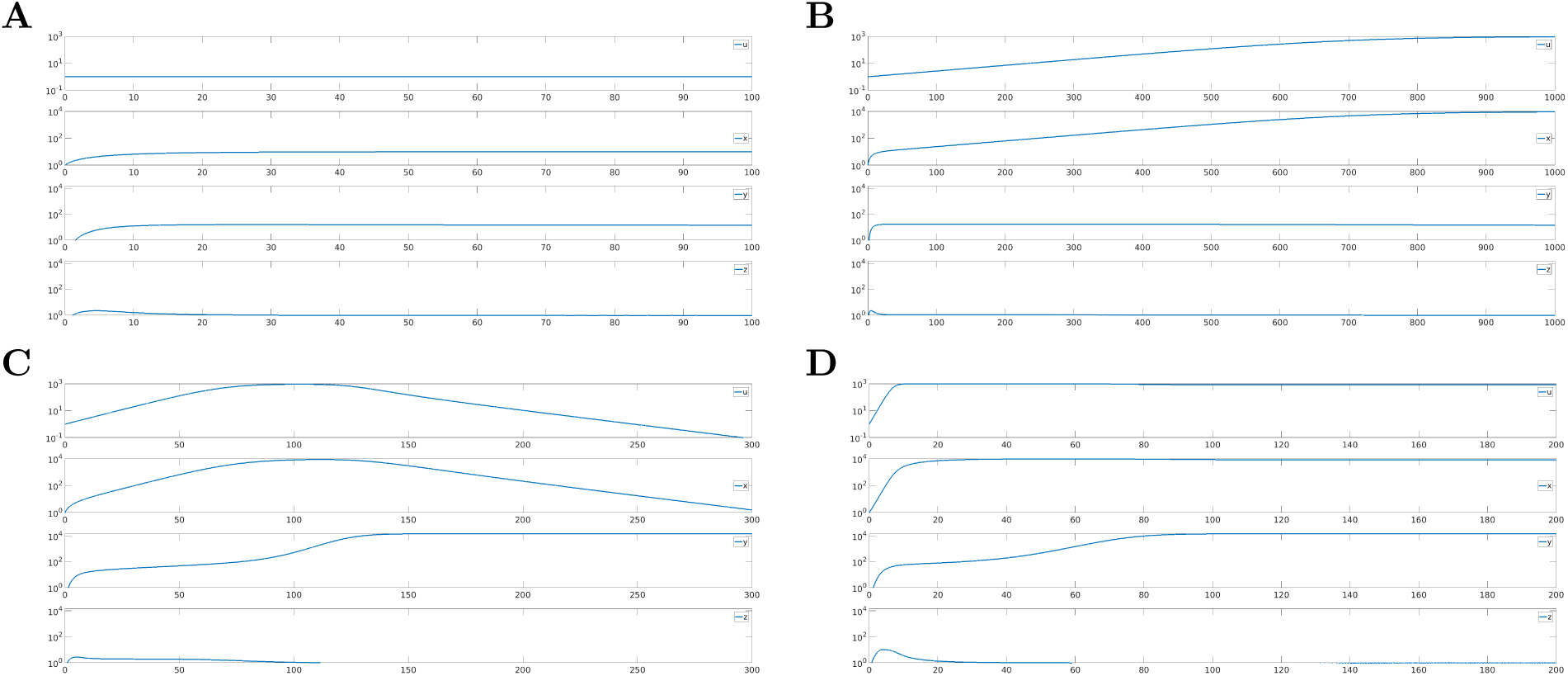
**A-D.** Simulations of system with “helper” intermediate, for λ = 10^-4^, 10^-2^, 10^-1^, and 1. (In **A**, *u*(*t*) converges to zero as *t* → œ, but very slowly.) Parameters as described in text for Figure 4. Initial states are always *u* = 1, × = 1, *y* = 0, *z* = 0. Only noticeable difference with simpler model is a slight delay in activation of *y*.

## 9 Degradation-based IFFL

The IFFL that we have been studying can be described phenomenologically by the diagram that is shown in the left panel of Figure 9: the input *u* activated both *x* and *y*, and *x* in turn represses the production of *y*. In the language of [2], this is a “type I” IFFL, a qualitative motif as shown in the center panel of Figure 9.

**Figure 9:**
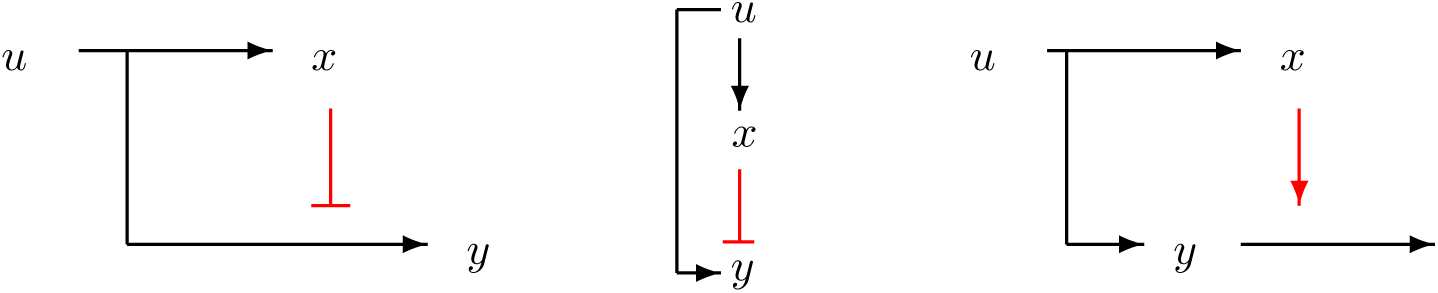
An incoherent feedforward loop (middle) and two instantiations, one through repression of production (left) and another one through enhancement of degradation (right). (Constitutive degradations of *x* and *y* not shown.)

An alternative phenomenological description of this same motif is in the right panel of Figure 9: now the regulatory node *x* serves to inactivate or degrade the output, rather than decreasing its formation. Thus, the same qualitative motif encompasses two very different molecular realizations, which may well differ significantly in their dynamic response and, ultimately, biological function.

A very simple set of ODE’s modeling the right-hand side motif is as follows:

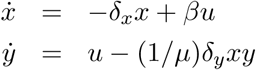

for suitable constants. It is easy to show that this system is also perfectly adapting. However, this second system does not exhibit scale invariance, see Figure 10.

**Figure 10:**
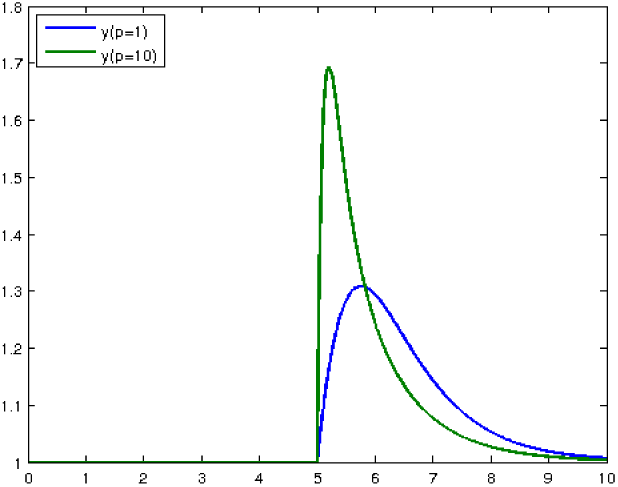
Different responses to *u*(*t*) and *pu*(*t*), of inactivation-based inhibition, showing failure of scale invariance. Input *u* switches from *u*(*t*) = 1 to *u*(*t*) = 2 at time *t* = 5. Parameters are *δ_x_* = *β* = *μ* = *δ_y_* = 1 and scale is *p* = 10.

Intuitively, multiplying *u* and *x* by *p* leads to a time-scale change in the output. This system is discussed as an example of a perfect adaptation in [63] (“sniffer”), and is analyzed in some detail in [57]. When the *y*-subsystem is fast compared to the first subsystem (*μ, δ_y_ ≫ δ_x_, β*), this system has an approximate scale-invariance property, because the output is approximately *u/x* after a short transient, and it appears generically in scale-invariant three-node enzymatic networks [54]. Further analysis showed, however, that there is an irreducible scale error, independent of how large the time scale gap is [53].

### Using a degradation-based IFFL in the model

We recall that the main model is as follows:

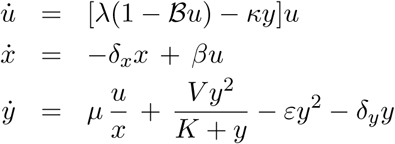

(including a carrying capacity term for convenience in numerical simulations). Suppose that we replace 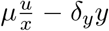 in the effector variable equation by *u − (1/μδ_y_xy*, as in the degradation-based IFFL. The full equations are now

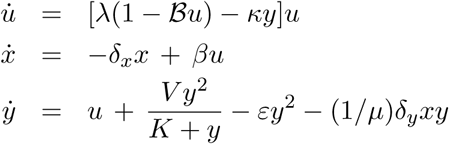

(again including a carrying capacity term). Using the same parameters as earlier, ***𝓑*** = 10^−3^, *V* = 0.25, *ε* = 10^−5^, *K* = 100, *κ* = 10^−5^, *μ* = 10, *d* = 0.1, the plots of solutions for various initial values of *u* (still in units of 10^6^ cells) are shown, for *t* ∊ [0,300], in Figure 11, as well as the familiar “two-zone” diagram is obtained at time *t* = 300, except that now we can see that the initial tumor load, keeping the rate λ constant, determines the outcome, at least at a fixed time.

**Figure 11:**
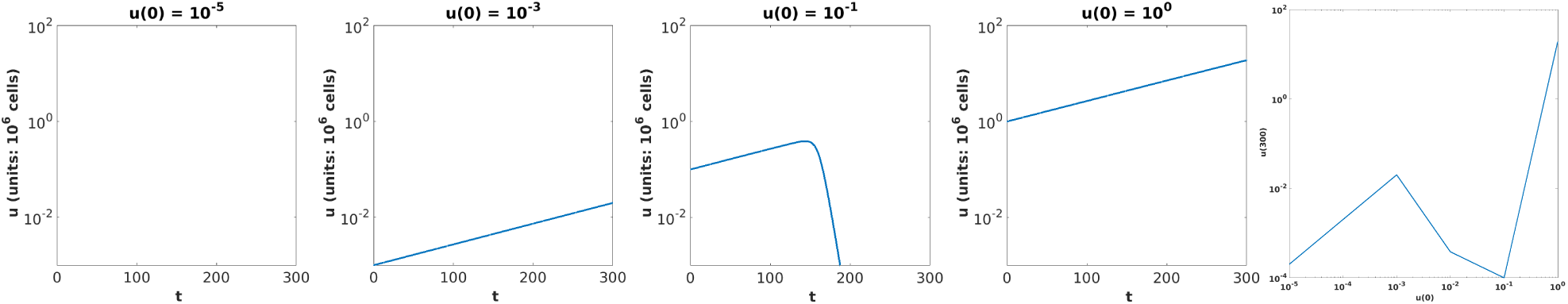
Simulations of degradation-IFFL model for various initial conditions, on interval *t* ∊ [0, 300] and with λ = 10^−2^. Right: Values of *u*(300) as a function of *u*(0), for a fixed value of λ = 10^−2^, in degradation-IFFL model (plot truncated below at 10^−4^). Parameters as in text.

## 10 Discussion and conclusions

The study of immune systems and their interactions with tumors has long been the focus of theoretical and mathematical immunology [28, 5]. Two influential contributions were the 1980 paper by Stepanova [61] in which a set of two ODE’s was used to represent tumor and immune system cells, and the 1994 paper by Kuznetsov, Makalkin, Taylor, and Perelson [38] in which a similarly simple model was used to provide an explanation for the sneaking-through phenomenon, though with escape of small tumors and with no mechanism for detection of rates of change of the immune challenge. It is impossible here to review the literature in this very active area of research; some reviews and textbooks are [5, 11, 3, 17, 69, 13, 65].

In this paper, we proposed a very simple phenomenological model that recapitulates some of the basic features of interactions between the immune systems and tumors (or more generally at this level of abstraction, other immune challenges) in the context of the estimation of tumor growth rates. The model leads to interesting conclusions regarding transitions between tolerance and elimination and the role of dynamics in self/nonself discrimination, and makes contact with several theory and experimental papers. Obviously, our model represents a purely phenomenological, macroscopic, and hugely over-simplified view of a highly complex, intricate, and still poorly understood network of interactions between different components of the immune system, as well as immune interactions with pathogens and tumors, Paraphrasing the well-known quote, our model is “as simple as possible but not simpler” to illustrate the particular phenomena of interest.

Since Paul Ehrlich’s work in 1909, [18] the interplay with the immune system has been a controversial, though recently accepted, aspect of cancer biology. These ideas were formalized in Burnet’s immune surveillance hypothesis [9], as an interaction between cancers that continuously arise and their repression by immune system, resulting in eventual elimination. Initial interest in this work was soon tempered by early experiments, but eventually new data led to a revival of these ideas in the late 1990s [15]. There is little doubt nowadays that immunosurveillance acts as a tumor suppressor, although it is also widely understood that the immune system can facilitate tumor progression by “sculpting” the immunogenic phenotype of tumors as they develop. Indeed, one current paradigm [16] is the so-called “three Es of cancer immunoediting” hypothesis: elimination, equilibrium, and escape. The first of this corresponds with the classical immunosurveillance idea: the immune system successfully eradicates the developing tumor. In the equilibrium or immune sculpting phase, the host immune system and any tumor cells that have survived the elimination phase enter into a dynamic quasi-steady state equilibrium during which the tumor cell population stays at sub-clinical levels. Ultimately, however, genetic and epigenetic heterogeneity in tumors, coupled by the Darwinian selection pressure exerted by the immune system, lead to the emergence of dominant clones with reduced immunogenicity which expand and become clinically detectable, and this is termed the escape phase. Our model does not directly address the effect of genetic or epigenetic modifications, and expanding it to do so remains a most interesting direction for further work.

Clinically, the interactions between the immune system and tumors are the focus of much current research because of the promise of novel immunotherapies such as checkpoint inhibitors [47]. It is worth pointing out the role of dynamical responses in immunotherapies, compared to classical chemotherapy and pathway inhibitors, which is emphasized in the UpToDate physician reference guide, from which we quote from the 01 Sep 2015 version: “patients may have a transient worsening of disease, manifested either by progression of known lesions or the appearance of new lesions, before disease stabilizes or tumor regresses.” (Interestingly, the solution in Figure 4B has this behavior.) This statement helps justify, in our view, the introduction of dynamical systems concepts into the field of immunotherapy.

